# Candidate genes-based investigation of susceptibility to Human African Trypanosomiasis in Côte d’Ivoire

**DOI:** 10.1101/186718

**Authors:** Bernardin Ahouty, Mathurin Koffi, Hamidou Ilboudo, Gustave Simo, Enock Matovu, Julius Mulindwa, Christiane Hertz-Fowler, Bruno Bucheton, Issa Sidibé, Vincent Jammoneau, Annette MacLeod, Harry Noyes, Simon-Pierre N’Guetta, for the TrypanoGEN Research Group as members of The H3Africa Consortium.

## Abstract

Human African Trypanosomiasis (HAT) or sleeping sickness is a Neglected Tropical Disease. Long regarded as an invariably fatal disease, there is increasing evidence that infection by *T. b. gambiense* can result in a wide range of clinical outcomes, including latent infections, which are long lasting infections with no parasites detectable by microscopy. The determinants of this clinical diversity are not well understood but could be due in part to parasite or host genetic diversity in multiple genes, or their interactions. A candidate gene association study was conducted in Côte d’Ivoire using a case-control design which included a total of 233 subjects (100 active HAT cases, 100 controls and 33 latent infections). All three possible pairwise comparisons between the three phenotypes were tested using 96 SNPs in16 candidate genes (*IL1, IL4, IL4R, IL6, IL8, IL10, IL12, IL12R, TNFA, INFG, MIF, APOL1, HPR, CFH, HLA-A* and *HLA-G*). Data from 77 SNPs passed quality control. There were suggestive associations at three loci in *IL6* and *TNFA* in the comparison between active cases and controls, one SNP in each of *APOL1*, *MIF* and *IL6* in the comparison between latent infections and active cases and seven SNP in *IL4, HLA-G* and *TNFA* between latent infections and controls. No associations remained significant after Bonferroni correction, but the Benjamini Hochberg false discovery rate test indicated that there were strong probabilities that at least some of the associations were genuine.

The excess of associations with latent infections despite the small number of samples available suggests that these subjects form a distinct genetic cluster different from active HAT cases and controls, although no clustering by phenotype was observed by principle component analysis. This underlines the complexity of the interactions existing between host genetic polymorphisms and parasite diversity.

**Author summary:** Since it was first identified, human African trypanosomiasis (HAT) or sleeping sickness has been described as invariably fatal. Recent data however suggest that infection by *T. b. gambiense* can result in a wide range of clinical outcomes in its human host including long lasting infections, that can be detected by the presence of antibodies, but in which parasites cannot be seen by microscopy; these cases are known as latent infections. While the factors determining this varied response have not been clearly characterized, the effectors of the immune responses have been partially implicated as key players. We collected samples from people with active HAT, latent infections and controls in endemic foci in the Côte d’Ivoire. We tested the role of single nucleotide polymorphisms (SNPs) in 16 genes on susceptibility/resistance to HAT by means of a candidate gene association study. There was some evidence that variants of the genes for *IL4, IL6, APOL1, HLAG, MIF* and *TNFA* modified the risk of developing HAT. These proteins regulate the inflammatory response to many infections or are directly involved in killing the parasites. In this study, the results were statistically weak and would be inconclusive on their own, however other studies have also found associations in these genes, increasing the chance that the variants that we have identified play a genuine role in the response to trypanosome infection in Côte D’Ivoire.

## Introduction

Sleeping sickness, or human African Trypanosomiasis (HAT), is caused by *Trypanosoma brucei gambiense* and *T.b. rhodesiense* and is transmitted by tsetse flies (*Glossina spp*)*. T. b. gambiense* is associated with a more chronic disease that can take decades to become patent, and *T.b. rhodesiense* causes an acute disease within months of infection. The chronic form of the disease caused by the *T. b. gambiense* is classically characterized by an early hemolymphatic stage (stage 1) associated with non-specific symptoms such as intermittent fevers and headaches, followed by a meningo-encephalitic stage (stage 2) in which the parasite invades the central nervous system and causes neurological disorders and death if left untreated. This chronic form is found in Western and Central Africa while the acute form caused by *T. b. rhodesiense* is endemic to Eastern Africa [1,2]. However, recent observations are increasingly indicating that infection by *T. b. gambiense* can result in a wide range of clinical outcomes in its human host [3,4]. Self-cure processes have been described and reviewed in Checchi *et al.* [3]. Furthermore, some authors argued that HAT is not invariably fatal [5], supporting observations made by Garcia et *al*. [4] who followed individuals who remained seropositive for HAT but without detectable parasites for two years. This clinical diversity is not well understood but could be due to parasite genetic diversity [6,7], human immune gene variability [8] or their interaction [9]. It has been suggested that genetic polymorphisms of the parasite could be associated with asymptomatic and very chronic infections [10]. Nevertheless, genes involved in the host immune response have been implicated in the control of infection or susceptibility to HAT [11,12] and also *T. congolense* infections in experimental models [13,14]. Among the genes implicated in the pathogenesis of the disease, are *IL1, IL4, IL6, IL8, IL10, IL12, IFNG, TNFA*, *CFH* and *MIF* [8,11,15-17].

Some studies have found associations between certain polymorphisms in genes encoding cytokines. For example, *IL4* plays a role in susceptibility to *T. brucei* infection [18], while polymorphisms in the *IL6* and *IL10* genes have been associated with a decreased risk of developing HAT [11,15]. On the other hand, polymorphisms in *TNFA* genes have been associated with an increased risk of developing the disease [11,15]. Furthermore, MacLean et *al*. [19] reported in their study that plasma levels of *IFNG* significantly decrease during the late stage of the *T.b. rhodesiense* disease [19]. Human blood contains trypanolytic factors (TLF1 and TLF2) that are lytic to almost all African trypanosomes except *T. b. rhodesiense* and *T. b. gambiense* [20]. Human TLF1 contains two primate-specific proteins, apolipoprotein L1 (APOL1) and haptoglobin-related protein (HPR). *APOL1* expression is induced by *T. b. gambiense* infection but expression is not associated with susceptibility to sleeping sickness [21]. Macrophage migration inhibitory factor (MIF) contributes to inflammation-associated pathology in the chronic phase of *T. brucei* infection [16]. Although genes directly involved in the immune response, like genes encoding cytokines, are very important candidates, genes implicated in the regulation of immunity also have a critical role. Thus, Courtin et *al*. [12] have shown a genetic association between *HLA-G* polymorphisms and susceptibility to HAT.

There is now cumulative evidence that polymorphisms in genes involved in the control of the immune response and genes implicated in the regulation of immunity could play a role in HAT infection outcome [11,12].

We report a candidate gene association study of the role of single nucleotide polymorphisms (SNPs) in *IL1*, *IL4, IL4R, IL6, IL8, IL10, IL12, IL12R, TNFA, INFG, MIF, HPR, CFH, APOL1, HLA-A*, and *HLA-G* genes on susceptibility/resistance to HAT.

## Results

### Characteristics of the population

A total of 100 cases (or former cases), 100 controls and 33 latent infections were enrolled into the study (Table 1). The mean age (range) of the study population was 38.8 (6–84) years. The sex ratio (male: female) was 0.88 (109/124).

**Table 1:** Population characteristics. **Table 1:** Population characteristics. Cases: CATT+ve, microscopy +ve; Controls: CATT-ve and microscopy–ve. Trypanolysis negative (TL-ve) cases were from subjects who were retrospectively sampled for the purpose of this study and who had not been subject to the trypanolysis test when originally identified as HAT cases by CATT and microscopy. Latent infection: subjects who were CATT positive (CATTpl ≥1/4) but parasites could not be detected by microscopy at repeated tests for at least two years.

|  | Cases<br>(n=100) | Controls<br>(n=100) | Latent infections<br>(n= 33) | Whole population<br>(n=233) |
| --- | --- | --- | --- | --- |
| Age mean | 41,8 | 36,8 | 36 | 38,8 |
| Males | 53 | 42 | 14 | 109 |
| Females | 47 | 58 | 19 | 124 |
| TL+ | 65 | 0 | 18 | 83 |
| TL- | 35 | 100 | 15 | 150 |

### Association study

Three association analyses were run on the three possible pairwise combinations of the controls, latent infections and active HAT cases. After filtering out SNP loci that were not in Hardy Weinberg equilibrium (P <0.001), or had > 10% missing genotypes or had minor allele frequencies of zero, 77 SNP loci remained in the latent infection vs active case comparison and 74 SNP loci remained in the other two comparisons. One individual was filtered out of the case sample and one out of the latent infection sample because of low genotyping rates (<90%).

FST between ethnic groups showed that allele frequency differences between the ethnic groups were small, indicating that ethnicity was unlikely to confound results (median Fst - 0.00011763, Fst maximum 0.015). Scatter plots of the first two principal components between the different phenotypes (cases, latent infections and controls) and different ethnic groups show that this population is homogeneous and that the samples did not cluster by phenotype or by ethnic groups (Fig 1) and consequently the data were not stratified by ethnicity.

**Fig 1.**
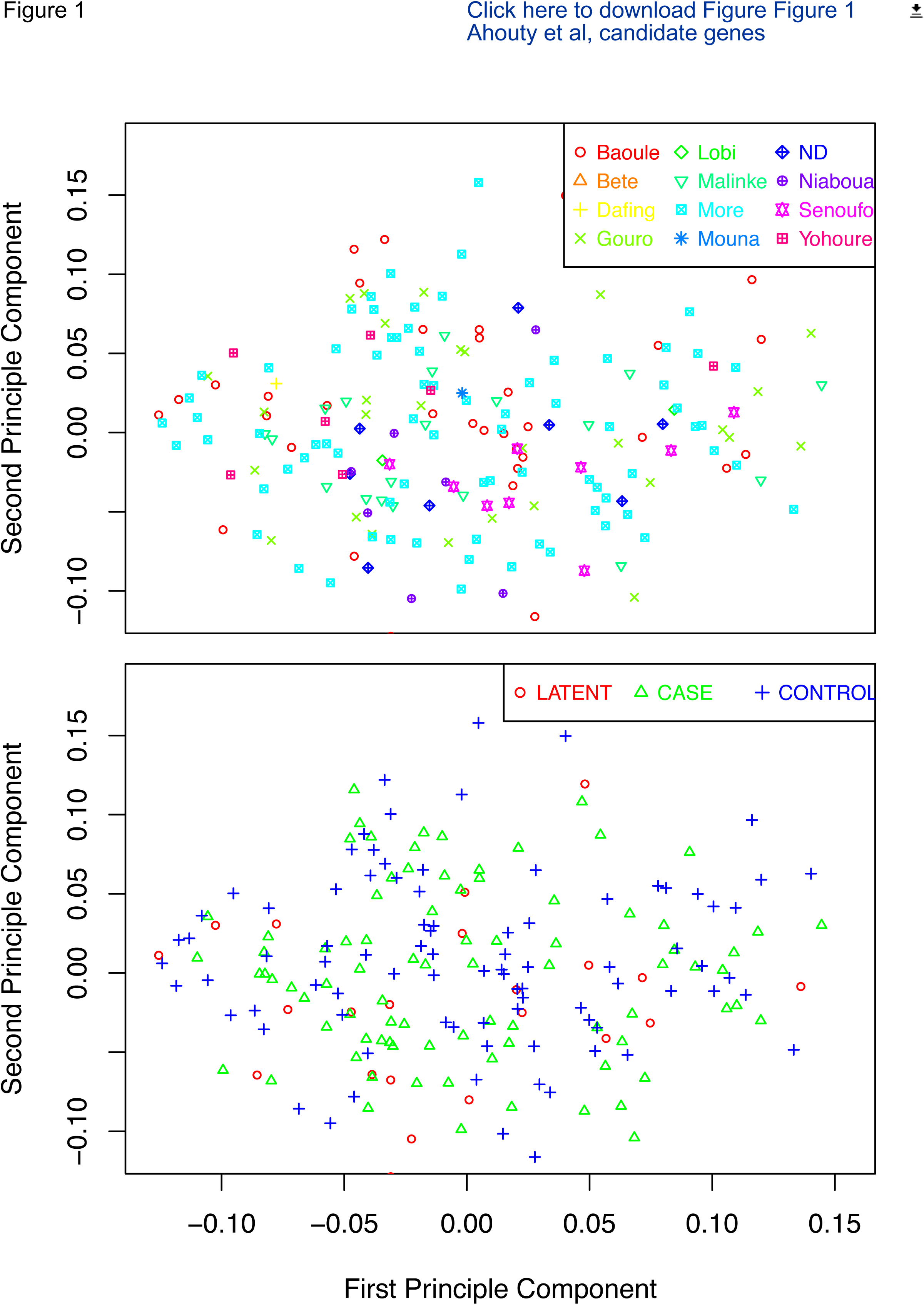
Multidimensional scaling (ms) plots of the genotype data by ethnicity (upper plot) and phenotype (lower plot). The plots show no evidence of clustering either by ethnicity or phenotype.

There were suggestive associations at three loci in *IL6* and *TNFA* in the comparison between active cases and controls (Table 2), three loci in *APOL1*, *MIF* and *IL6* in the comparison between latent infections and active cases (Table 3) and five loci in *IL4* (Fig 2), and one each in *HLA-G* and *TNFA* between latent infections and controls (Table 4). After Bonferroni correction, none of these associations remained significant, however Bonferroni correction is very conservative, particularly since there was some linkage between adjacent marker SNP in some genes. The Benjamini-Hochberg false discovery rate (FDR) test, which shows the probability that an observation is a false positive, indicated that at least some of the suggestively positive samples may be true positives. Under the FDR the rs62449495 SNP in *IL6* had an 80% chance of being a true positive and the *TNFA*-308 rs1800629 SNP had a 71% chance of being a true positive (Table 2). The strongest association was with *APOL1* rs73885319, which is also known as the G1 allele, which had a 90% chance of being a true positive in the comparison between latent infections and controls (Table 3). Complete results for all tests that passed quality control are shown in supplementary data tables S1, S2 and S3.

**Fig 2.**
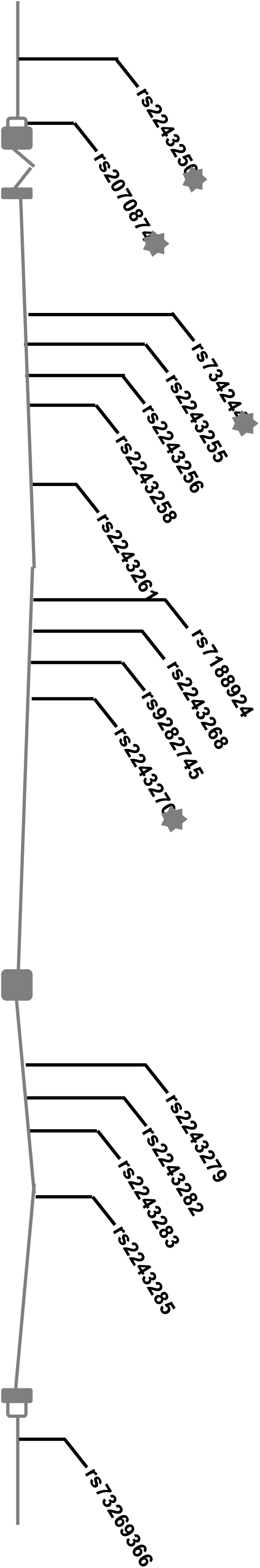
Positions of SNP genotyped within IL4. SNP with suggestive associations (uncorrected p <0.05) are indicated with a star.

**Table 2:**
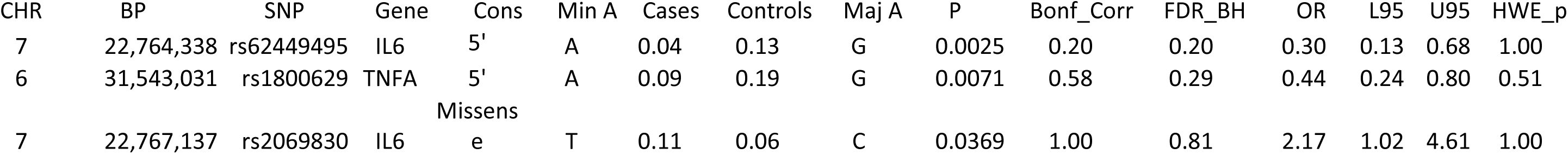
Association tests between cases (n=99) and controls (n=100)

**Table 3:**
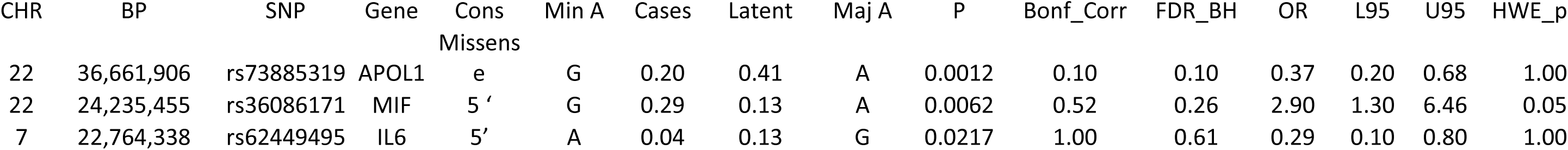
Association tests between latent infections (n=32) and active HAT cases (n=99)

**Table 4:** Association tests between latent infections (n=32) and controls (n=100)

| CHR | BP | SNP | Gene | Cons | Min A | Latent | Controls | Maj A | P | Bonf_Corr | FDR_BH | OR | L95 | U95 | HWE_p |
| --- | --- | --- | --- | --- | --- | --- | --- | --- | --- | --- | --- | --- | --- | --- | --- |
| 5 | 132,009,154 | rs2243250 | IL4 | 5' | C | 0.42 | 0.27 | T | 0.0159 | 1.00 | 0.48 | 2.02 | 1.13 | 3.64 | 0.80 |
| 5 | 132,010,726 | rs734244 | IL4 | Intronic | T | 0.25 | 0.41 | C | 0.0215 | 1.00 | 0.48 | 0.49 | 0.26 | 0.92 | 0.01 |
| 5 | 132,016,554 | rs2243282 | IL4 | Intronic | A | 0.30 | 0.45 | C | 0.0347 | 1.00 | 0.48 | 0.53 | 0.29 | 0.96 | 1.00 |
| 6 | 29,799,116 | rs1611139 | HLA-G | 3' | T | 0.26 | 0.14 | G | 0.0354 | 1.00 | 0.48 | 2.13 | 1.04 | 4.36 | 0.39 |
| 5 | 132,009,710 | rs2070874 | IL4 | 5' UTR | T | 0.34 | 0.50 | C | 0.0372 | 1.00 | 0.48 | 0.53 | 0.30 | 0.96 | 0.69 |
| 6 | 31,543,031 | rs1800629 | TNFA | 5' | A | 0.08 | 0.19 | G | 0.0396 | 1.00 | 0.48 | 0.37 | 0.14 | 0.99 | 0.51 |
| 5 | 132,014,109 | rs2243270 | IL4 | Intronic | A | 0.36 | 0.23 | G | 0.0415 | 1.00 | 0.48 | 1.88 | 1.02 | 3.45 | 1.00 |

Cons= Consequence from Ensembl Variant predictor, Min A = minor allele, Maj A = major allele, Latent minor allele frequency of Latent cases; Controls minor allele frequency of controls; OR = odds ratio, P = p-value, [L95-U95] = confidence interval of odds ratio, Bonf\_Corr= Bonferroni corrected p value, FDR\_BH = False Discovery Rate Benjamini-Hochberg (it is probability of falsely rejecting the null hypothesis. The null hypothesis is that allele frequencies in cases and controls are the same), OR = odds ratio, [L95-U95] = confidence interval; HWE P is the Hardy- Weinberg p-value.

## DISCUSSION

In this study, we investigated the role of single nucleotide polymorphisms on susceptibility/resistance to HAT. We genotyped 96 SNPs within sixteen genes that were investigated to test genetic association with HAT in Côte d’Ivoire. For consistency, all samples were commercially genotyped using two platforms: Genome Transcriptome de Bordeaux, France and LGC Genomics UK.

Our results suggest that three SNPs were associated with the development of HAT by controls, another three were associated with the development of active HAT by people with latent infections, and seven SNP were associated with the risk of developing latent infections five of them in *IL4* although none of these results remained significant after Bonferroni correction. Only two of the SNP with suggestive associations were coding and missense (Il6 rs2069830 and APOL1 rs73885319) since, with some exceptions, SNP were primarily selected as tags that might be linked to functional SNP rather than for any putative function. The most striking feature of the results is that most associations were found in the comparisons with latent infections, despite limited power, with only 33 of these samples being available compared with 100 each for the cases and controls. This illustrates the increased power to be gained from using well defined phenotypes. Given the evidence that some people in West Africa can self-cure their *T. b. gambiense* infections [5] it is likely that some of the control subjects are resistant to infection or have recovered from infection, whilst others are naïve and susceptible. These two groups may have very different genetic backgrounds which could confound the association studies. The data also suggest that the latent infections may represent a genetically distinctive group. Although the principle component analysis did not identify any cluster associated with latent infections (Fig 1) further studies of their genetic and immunological profiles are required to test this hypothesis.

### Associations with *IL6*

The minor (A) allele of *IL6* rs62449495 appeared to protect against progression from latent infections to active HAT (Table 3) and against the development of active HAT by controls but these associations did not remain significant after Bonferroni correction (Table 2). The major allele of rs2069830 was also protective against the development of active HAT by controls before but not after Bonferroni correction (Table 2). The rs2069830 SNP causes a proline to serine change, but this is predicted to be benign by both Sift and Polyphen according to the Ensembl Variant Effect predictor. IL6 plays a key role in the acute inflammatory response and in regulation of the production of acute phase proteins such as C-reactive protein [22]. It contributes to the inflammatory response, regulates haematopoiesis, which may contribute to the anaemia associated with HAT, and modifies the permeability of the blood brain barrier, which may contribute to the development of the stage 2 invasion of the CNS [22,23]. The results of our association study are consistent with the involvement of *IL6* variants in the susceptibility to the disease as Courtin *et al* [15] have reported before. However, it should be noted that the polymorphisms of *IL6* in our study (rs62449495 and rs2069830) are different of those found by Courtin *et al.* [15] with *IL6*4339= rs2069849. They showed that in DRC, *IL6*4339 SNP was significantly associated with a decreased risk of developing the disease with a *P*-value = 0.0006 before the Bonferroni correction (0.04 after Bonferroni Correction).

### Association with *TNF A*

Our results also indicate that subjects carrying the A allele of *TNFA* (rs1800629) A/G had a lower risk of developing active HAT (Table 2) or a latent infection (Table 4), suggesting the possibility of a protective effect. This SNP is also known as the *TNF*-308 SNP and the minor A allele is associated with higher plasma levels of *TNFA* [24]. In a previous study in Côte d’Ivoire, Courtin et *al*. [11] showed that the distribution of the *TNFA-*308G/A polymorphism did not differ significantly between cases and controls in the total population, but found that, under a recessive model the AA genotype was associated with risk of HAT in the 39 cases and 57 controls who had been living in the endemic area of Sinfra for less than 10 years (before Bonferroni correction). In contrast, in our study, all participants had lived in the endemic area all their lives and the association was additive rather than recessive, furthermore the MAF in Courtin’s studies was 24%, whilst in ours it was 14%, suggesting significant differences in population structure between the two studies. Varying results in *TNFA* associations studies are not uncommon, multiple candidate gene association studies of *TNFA-*308 (rs1800629) and malaria have found inconsistent results in different populations [25,26]. Another explanation could be that the *TNFA-*308 has no effect but is in linkage disequilibrium (LD) with another unidentified polymorphism. Varying LD across populations might lead to different findings [27]. Despite the conflicting results from association study, animal models indicate that *TNFA* is likely to be a key mediator in the control of *T. brucei* infections [28] and a direct dose dependent lytic effect of *TNFA* on purified *T.b. gambiense* parasites has been reported suggesting an involvement in parasite growth control [29,30]. Given the experimental evidence for a role for TNFA, it is possible that inconsistent results of association studies are a consequence of different *TNFA* alleles only making a small difference to infection outcome and that therefore larger studies would be needed to detect associations and also to heterogeneity in regulation of *TNFA* between populations.

### Association with *IL4*

Five of the seven SNP that were suggestively associated with the comparison between latent infections and controls were in *IL4* (Table 4, Fig 2). Three of these SNP were adjacent to each other at the 5’ end of the gene (Fig 2), although linkage between them was modest (r^2^ <0.5). While the individual associations did not remain significant after Bonferroni correction the observation that five of the sixteen SNP tested in *IL4* had suggestive associations increases the probability of a genuine association with *IL4* and the risk of developing a latent infection. A study using three *IL4* SNP in DRC did not find any associations, but that was in a comparison between cases and controls [15]. In B10.Q mice deletion in *IL4* lead to increased *T. brucei* parasitaemia levels but longer survival time [18]. The relationship between *IL4* polymorphisms and acute HAT infection requires further study.

### Association with *APOL1*

The suggestive association with *APOL1* G1 allele rs73885319 and protection against progression from latent infection to active HAT is consistent with the association found in Guinea [31,32], although in those studies this SNP was also associated with increased risk of controls developing a latent infection as well. Moreover, *APOL1* expression is induced by *T. b. gambiense* infection but not associated with differential susceptibility to sleeping sickness [21]. *APOL1* rs73885319 is also known as the G1 allele of *APOL1* and is associated with kidney disease in African Americans and the relatively high frequency of this deleterious allele was assumed to be due to selection by HAT [33]. The data presented here is consistent with that hypothesis, but no support was found for the role of the G2 allele of *APOL1* (rs71785313) in HAT although it is implicated in kidney disease.

### Association with *MIF*

Although, our data show that subjects carrying the G allele of *MIF* (rs36086171) G/A had a risk of developing active HAT (Table 3), we did not find a significant difference after correction (BONF = 0.52). Macrophage inhibitory factor (MIF) is a ubiquitously expressed protein that has proinflammatory, hormonal and enzymatic activities [34]. It is implicated in many inflammatory diseases [35]. It functions by recruiting myeloid cells to the site of inflammation [36], by inducing their differentiation towards M1 cells secreting TNF [37] and by suppressing p53-dependent apoptosis of inflammatory cells. While there are no human studies directly linking *MIF* to HAT, murine studies show that *MIF* plays a role as a mediator of the inflammation which is a key feature in trypanosomiasis-associated pathology [16,38].

### Association with *HLA-G*

*HLA-G* molecule plays an important role on immune response regulation and has been associated with the risk of human immunodeficiency virus (HIV) infection [39], human papilloma virus [40] and herpes simplex virus type 1 [41]. The results of our association study are consistent with the involvement of HLA-G genetic variants in the susceptibility to the disease [12]. The rs1611139 T allele of the *HLA-G* gene showed a suggestive association with increased risk of controls a latent infection. This result could be due to differences in selection pressures possibly driven by variability in the immuno-pathology of the diseases. It is known that cytokines such as IL-10 can induce HLA-G expression by affecting mRNA transcripts and protein synthesis by human monocytes and trophoblasts, thus having a significant impact on parasitic infections [42].

## Conclusion

The results discussed in this paper should be used with caution as no loci remained significantly associated with HAT after the Bonferroni correction for multiple testing, although some had high probabilities of being associated using a false discovery rate test. Some of the SNP loci identified here were also significant in other studies. Multiple independent observations of marginally significant effects suggest that these effects may be genuine but that the effect size is not large enough to be detected by the numbers available to be tested.

Our data support the findings from Guinea about the role of the *APOL1* G1 allele and also suggest that the polymorphisms of *IL4, IL6, HLA-G* and *TNFA* present interesting candidates for the investigation of the genetic susceptibility / resistance to HAT.

The large number of suggestive associations with latent infections despite the small number of samples indicate the people with latent infections form a genetically distinctive group that merit further investigation.

## Material and Methods

### Population study, definition of phenotypes and study design

The study took place in western-central Côte d’Ivoire in the main HAT foci of Bonon, Bouafle, Zoukougbeu, Oume, and Sinfra. Samples were collected in three stages, (i) previously archived samples; existing collections of previously archived samples were centralized to CIRDES (Centre International de Recherche-Développement sur l’Elevage en zone Subhumide) Burkina-Faso. (ii) retrospectively collected samples; study sites were revisited to consent and resample previously diagnosed and treated patients and (iii) prospectively collected samples including new HAT patients. Three phenotypes were considered: (i) cases, defined as individuals in whom the presence of trypanosomes was confirmed by microscopy; (ii) controls, defined as individuals living in the endemic area who are card agglutination test for trypanosomiasis (CATT) negative and trypanolysis test (TL) negative, and without evidence of previous HAT infection and (iii) latent infections, defined as subjects who were CATT positive (CATTpl ≥1/4) but in whom parasites could not be detected by microscopy at repeated tests for at least two years.

A total of 233 individuals with 100 former HAT cases, 100 controls and 33 latent infections were included in the study. Samples were frozen directly in the field at −20°C and kept at that temperature until use. For each individual, an aliquot of plasma was used to perform the immune trypanolysis test that detects Litat 1.3 and Litat 1.5 variable surface antigens specific for *T.b. gambiense* [43]. All control individuals (n=100) included in this study were negative on the trypanolysis test (TL-ve) (Table 1).

This study was one of five studies of populations of HAT endemic areas in Cameroon, Côte d’Ivoire, Guinea, Malawi and Uganda by the TrypanoGEN consortium [44]. The studies were designed to have 80% power to detect odds ratios (OR) >2 for loci with disease allele frequencies of 0.15 – 0.65 and 100 cases and 100 controls with the 96 SNPs genotyped. Power calculations were undertaken using the genetics analysis package gap in R [45].

### Selection of SNPs and genotyping

Genomic DNA was obtained from peripheral blood samples. Extraction was performed using the Qiagen DNA extraction kit (QIAamp DNA Blood Midi Kit) according to the manufacturer’s instructions and quantified by Nanodrop assay. DNA was stored at −20°C until analysis.

Ninety-six SNPs were genotyped in *IL1*, *IL4, IL4R, IL6, IL8, IL10, IL12, IL12R, TNFA, INFG, MIF, HPR, CFH, APOL1, HLA-A*, and *HLA-G*. The SNPs were selected by two strategies: 1) SNP in *IL4, IL6, IL8, HLAG* and *IFNG* were designed as markers for linkage disequilibrium (LD) scans of each gene [46], 2) SNPs in other genes were selected based on reports in the literature that they were associated with trypanosomiasis or other infectious diseases. The LD scans were designed using 1000 Genomes Project data [47], merged with low fold coverage (8-10x) whole genome shotgun data generated from 230 residents living in regions (DRC, Guinea Conakry, Côte D’Ivoire and Uganda) where trypanosomiasis is endemic (TrypanoGEN consortium, European Nucleotide Archive study EGAS00001002482). Loci with minor allele frequency < 5% in the reference data were excluded and an r^2^ of 0.5 was used to select SNP in linkage disequilibrium. DNA was genotyped by two commercial service providers: INRA-Site de Pierroton, Plateforme Genome Transcriptome de Bordeaux, France and ii-LGC genomics Hoddesden UK. At INRA multiplex design (two sets of 40 SNPs) was performed using Assay Design Suite v2.0 (Agena Biosciences). SNP genotyping was achieved with the iPLEX Gold genotyping kit (Agena Biosciences) for the Mass Array iPLEX genotyping assay, following the manufacturer’s instructions. Products were detected on a Mass Array mass spectrophotometer and data were acquired in real time with Mass Array RT software (Agena Biosciences). SNP clustering and validation was carried out with Typer 4.0 software (Agena Biosciences). At LGC Genomics SNP were genotyped using the PCR based KASP assay[48].

### Statistical analysis

All statistical analyses were performed using the Plink 1.9 and R v 3.2.1 software. Individuals were excluded who had > 10% missing SNP data. SNP loci were excluded that > 10% missing genotypes or if the control samples were not in Hardy-Weinberg equilibrium (p<0.001). Case-control association analysis using SNP alleles/genotypes was undertaken using Fisher’s exact test. The difference between the ethnic groups was estimated using the FST, with values potentially varying from zero (no population differentiation) to one (complete differentiation) [47]. Ethnicity was not used as a risk factor for HAT because we observed no significant differences in FST between ethnic groups (FST maximum 0.015) or clustering by linguistic group by multidimensional scaling in Plink. Some studies have used ethnicity and age as risk factors for HAT but found no significant association [11,15]. 77 SNP remained in at least one comparison after filtering. P values were adjusted for multiple testing using the Bonferroni corrections and Benjamini Hocheberg false discovery rate test as implemented in Plink.

### Ethics Statement

The population of the study was informed. All adult subjects provided written informed consent. For children, a parent or guardian of any child participant provided written informed consent on their behalf. The protocol of the study was approved by the traditional authorities (chief and village committee) and by National ethics committee hosted by the Public Health Ministry of Côte d’Ivoire with the number: N°38/MSLS/CNERm-dkn 5th May 2014. This study is part of the TrypanoGen project which aims to a better understanding of genetic determinism of human susceptibility to HAT and the TrypanoGen-CI samples were archived in the TrypanoGen Biobank hosted by CIRDES in Bobo-Dioulasso, Burkina Faso [44].

## Acknowledgements

We thank subjects who generously donated their specimens and the field workers from the Ivorian HAT foci for their dedication in collecting and processing these specimens.

